# Protein degradation in a TX-TL cell-free expression system using ClpXP protease

**DOI:** 10.1101/019695

**Authors:** Zachary Z. Sun, Jongmin Kim, Vipul Singhal, Richard M. Murray

## Introduction

An *in vitro* S30-based *Escherichia coli* expression system (“Transcription-Translation”, or “TX-TL”) has been developed as an alternative prototyping environment to the cell for synthetic circuits [1-5]. Basic circuit elements, such as switches and cascades, have been shown to function in TX-TL, as well as bacteriophage assembly [2, 6]. Circuits can also be prototyped from basic parts within 8 hours, avoiding cloning and transformation steps [7]. However, most published results have been obtained in a “batch mode” reaction, where factors that play an important role for *in vivo* circuit dynamics – namely protein degradation and protein dilution – are severely hindered or are not present. This limits the complexity of circuits built in TX-TL without steady-state or continuous-flow solutions [8-10]. However, alternate methods that enable dilution either require extra equipment and expertise or demand lower reaction throughput.

We explored the possibility of supplementing TX-TL with ClpXP, an AAA+ protease pair that selectively degrades tagged proteins [11], to provide finely-tuned degradation. The mechanism of ClpXP degradation has been extensively studied both *in vitro* and *in vivo* [12-15]. However, it has not been characterized for use in synthetic circuits – metrics such as toxicity, ATP usage, degradation variation over time, and cellular loading need to be determined. In particular, TX-TL in batch mode is known to be resource limited [16], and ClpXP is known to require significant amounts of ATP to unfold different protein targets [17, 18]. We find that ClpXP’s protein degradation dynamics is dependent on protein identity, but can be determined experimentally. Degradation follows Michaels-Menten kinetics, and can be fine tuned by ClpX or ClpP concentration. Added purified ClpX is also not toxic to TX-TL reactions. Therefore, ClpXP provides a controllable way to introduce protein degradation and dynamics into synthetic circuits in TX-TL.

## RESULTS AND DISCUSSION

There are two methods to supplement ClpXP into TX-TL – by expressing the proteins off of plasmid DNA or by adding purified protein. We first attempted to express both ClpX and ClpP off of DNA, and found that ClpX when expressed was able to degrade fluorescent ssrA-tagged reporters at a rate independent of ClpP concentration (data not shown). However, expressing ClpX posed two difficulties: degradation was rate-limited and time-delayed based on production rate of ClpX, and the load of ClpX expression reduced the expression of other synthetic circuits. To avoid these problems, we tried to purify a N-terminal His-tagged version of ClpX using a standard Ni-NTA procedure; however, we found that this version of ClpX lost activity after the purification procedure when stored in a previously developed glycerol-free buffer compatible with TX-TL [7]. We hypothesized that a linked-hexametric form previously developed would be more stable in solution [13]. With the His-tagged linked-hexameric N-terminal deletion form of ClpX, we were able to retain activity (**Fig. 1ab**). We refer to this as the ClpX form hereafter. To quantify degradation rates, we also purified fluorescent proteins with and without –ssrA tags (**Fig. 1c**).

**Figure 1.**
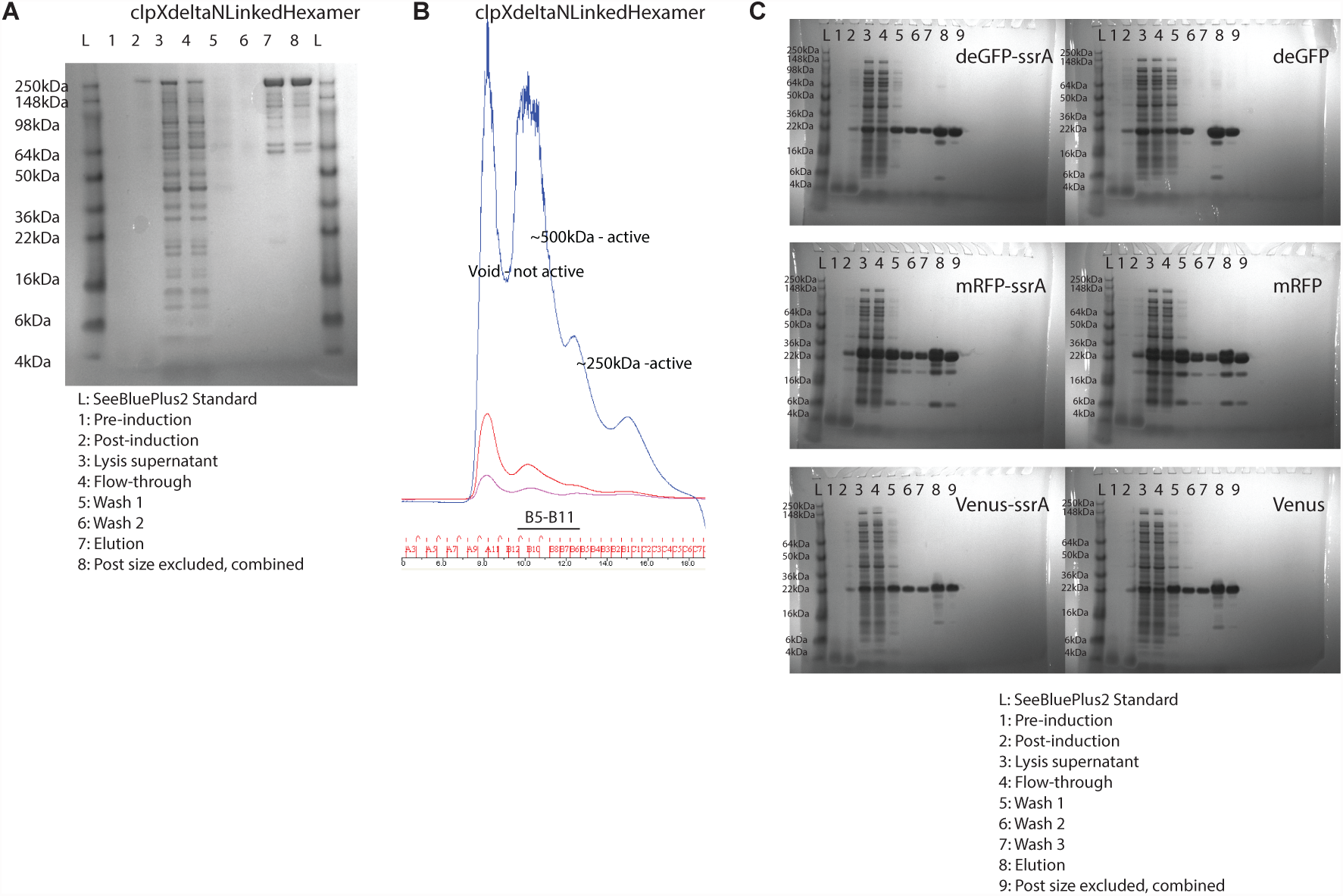
Purification of clpXdeltaNLinkedHexamer (“ClpX”) and fluorescent proteins. a) ClpX with a His6 tag is purified in a Ni-NTA column and run on denaturing SDS-PAGE gel. b) Size excluded chromatography of elute product on a Superdex 200 10/300 column, with a functionality test of active species. B5-B11 are fractions combined for forward use. c) Three fluorescent proteins (deGFP, mRFP, Venus), with or without ssrA tags, with a His6 tag are purified in a Ni-NTA column and run on denaturing SDS-PAGE gel. Last column is active, purified fractions combined for forward use.

We first verified the ability of purified ClpX to selectively degrade ssrA-tagged versions of deGFP, mRFP, and Venus over non-tagged versions (**Fig. 2**). ClpX is selective for ssrA-tagged versions over non-ssrA tagged versions, and has a degradation rate dependent on protein identity. This is similar to findings *in vitro*, where ClpX is known to have different unfolding rates and ATP hydrolysis rates depending on the target difficulty [17, 19]. We also determined the concentration dependence of ClpX to degradation rate and throughput (**Fig. 3ab**). Initially, we made a simple Michelis-Menten model:

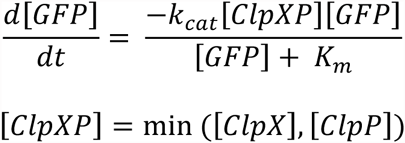

**Figure 2.**
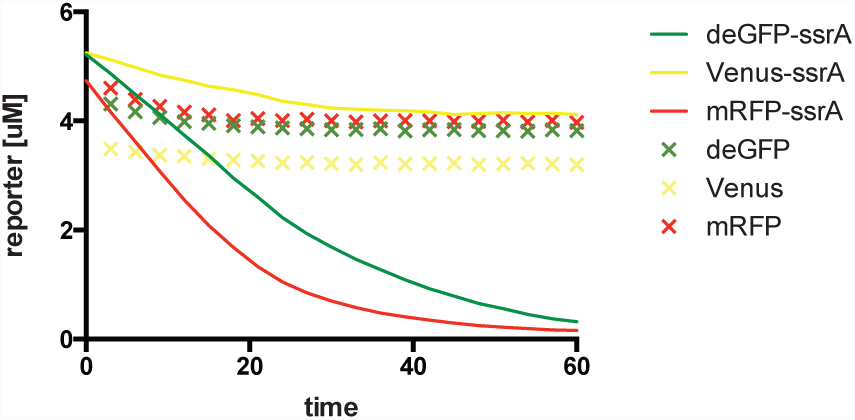
Degradation of deGFP, Venus, and mRFP by added ClpX. 400 nM of purified ClpX is combined with approximately 5 μM of purified fluorescent proteins with and without ssrA degradation tags in TX-TL.

**Figure 3.**
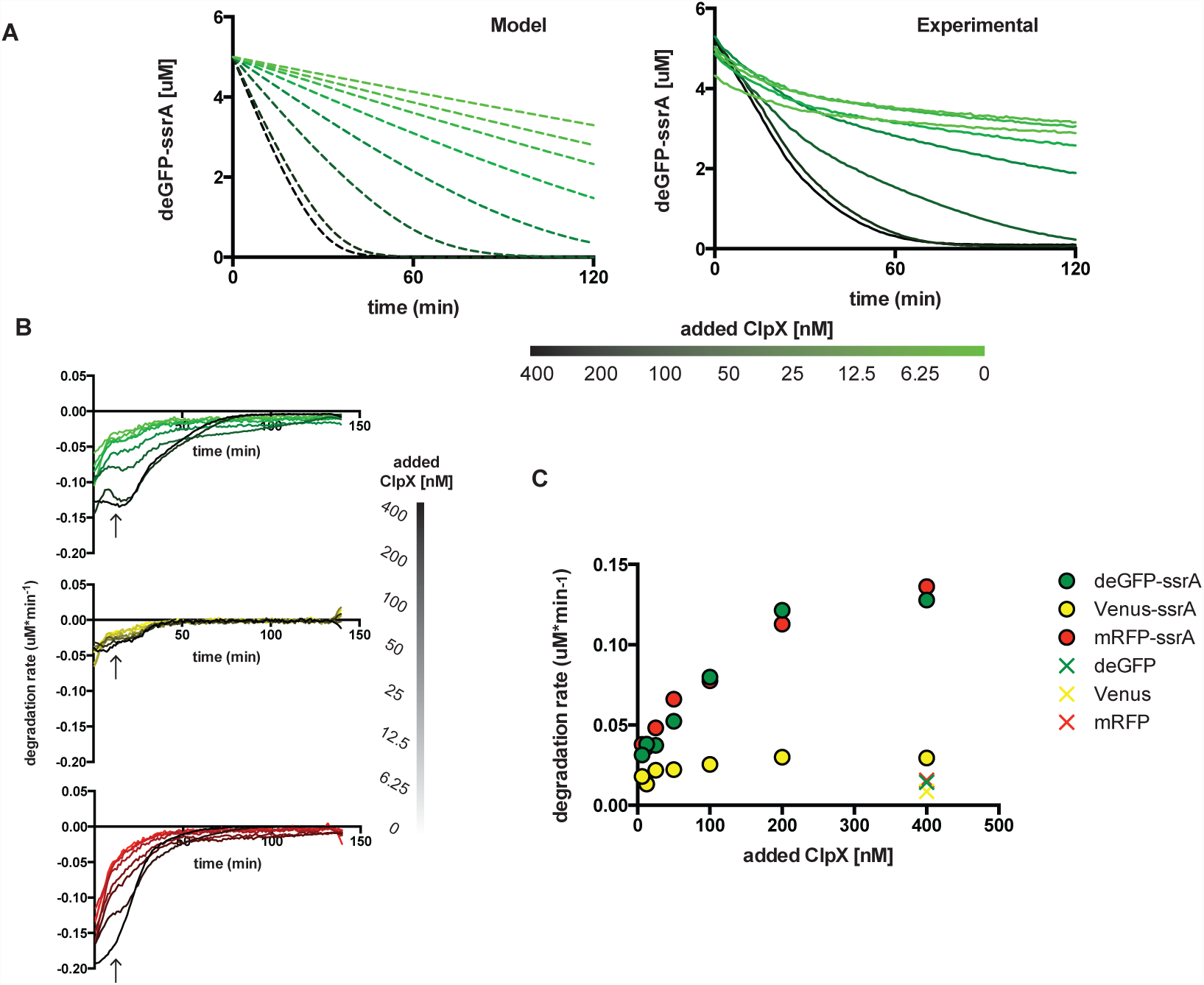
Dependence of degradation on ClpX concentration. A) Shown to the left is a Michaelis-Menten kinetics model of degradation of fluorescent GFP-ssrA over time, using K_m_ 1.1 μM, k_cat_ 0.9 min^-1^, similar to literature values [14]. Added ClpX is varied from 400nM to 0nM. Showing to the right is experimental data demonstrating similar results as modeling data for deGFP-ssrA. B) Degradation rate of deGFP-ssrA, Venus-ssrA, and mRFP-ssrA over time with varied ClpX concentration. At the arrow is t =16.5 min. C) Degradation rate plotted at t = 16.5 min as a function of added ClpX, for both ssrA tagged and non-ssrA tagged fluorescent proteins.

We set K_m_ to 1.1 μM, k_cat_ to 0.9 min^-1^ based on previously published parameters [14], and estimated initial (native) concentrations of 20 nM ClpX and of 250 nM ClpP in TX-TL extract. We also ran a TX-TL experiment varying added ClpX concentrations from 0 nM to 400 nM. Comparing the modeling data to the experimental data, ClpXP degradation closely follows Michaelis-Menten kinetics as described. We also determined concentration dependence of ClpX to Venus-ssrA and mRFP-ssrA, and then plotted the degradation rate vs. time **(Fig. 3c)**. Since the degradation rate changes over time as the concentration of substrate decreases, we plotted degradation rate vs. added ClpX concentration at t = 16.5 min, which we took as an initial degradation rate **(Fig. 3d)**. In this figure, the dependence of fluorescent reporter to added ClpX concentration can be clearly discerned, with a saturation point between 200-400 nM of additional ClpX added to the system, suggesting a rate-limiting native ClpP concentration. The decreased degradation rate of Venus-ssrA relative to deGFP-ssrA and mRFP-ssrA can also be seen. Based on this figure, additional ClpX is able to increase deGFP-ssrA degradation rates by 5-fold, mRFP-ssrA degradation rates by 4.2-fold, and Venus-ssrA degradation rates by 1.8-fold.

We additionally tried to supplement TX-TL with ATP and Mg, based on the known heavy usage of ClpX for protein unfolding (500 ATP per titin I27 subunit, ∼100 aa) [19]. With 200 nM of added ClpX in TX-TL, there was a clear effect on the degradation of Venus-ssrA over deGFP-ssrA and mRFP-ssrA, indicating that at working concentrations of proteins likely achievable in a typical TX-TL reaction ATP concentration was not rate-limiting except on hard-to-degrade proteins (**Fig. 4**).

**Figure 4.**
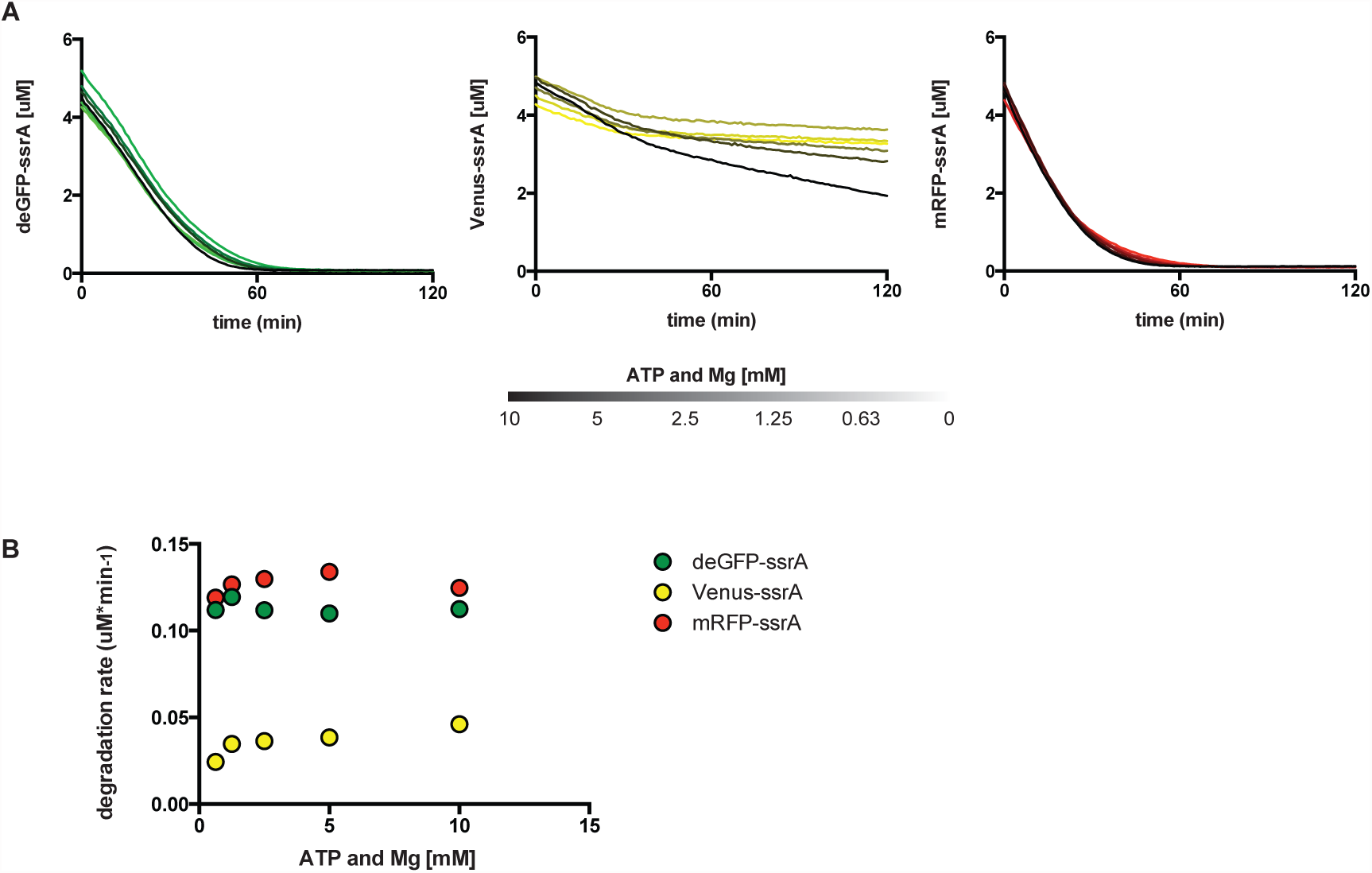
Dependence of degradation on added ATP and Mg. A) With added ClpX set to 200 nM, degradation of fluorescent ssrA tagged proteins is plotted as a function of time and additional ATP and Mg added to each reaction. B) Degradation rate at t = 16.5 min from above data is plotted as a function of additional ATP and Mg.

In order for supplemented ClpX to be a useful tool for controlled protein degradation, either the addition of ClpX or the ClpX storage buffer must not be toxic to expression of proteins off of DNA in TX-TL. We diagnosed this by diluting purified ClpX into a TX-TL reaction driving the production of deGFP off of a strong promoter, and saw no effect at saturating ClpX levels to DNA expression (**Fig. 5**). We also verified that ATP was not rate-limiting to ClpXP degradation and DNA expression by treating the degradation of mRFP-ssrA as a “background process” running during a TX-TL reaction producing deGFP (**Fig. 6**). As 10 μM of mRFP-ssrA is approaching the maximum protein concentration producible by TX-TL, this result suggests that ATP is at a saturating level in a typical TX-TL reaction. However, we did not test the combination of a difficult-to-degrade protein such as Venus-ssrA combined with the expression of saturating amounts of mRFP, which would support this conclusion.

**Figure 5.**
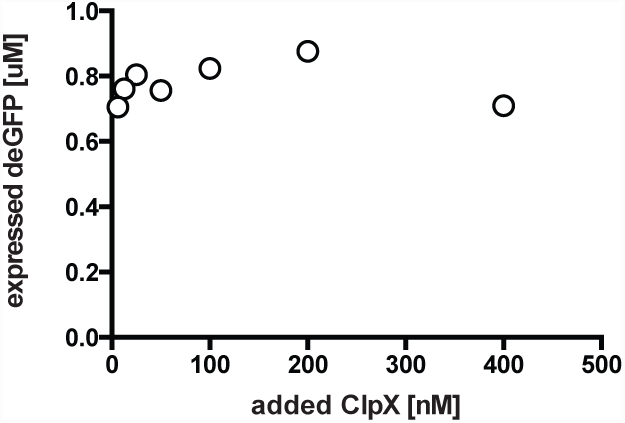
Toxicity of added ClpX to TX-TL expression. A plasmid expressing deGFP off of a strong promoter-UTR is added to a TX-TL reaction at 1 nM, and endpoint expression after 8 hours is plotted as a function of added, purified ClpX protein.

**Figure 6.**
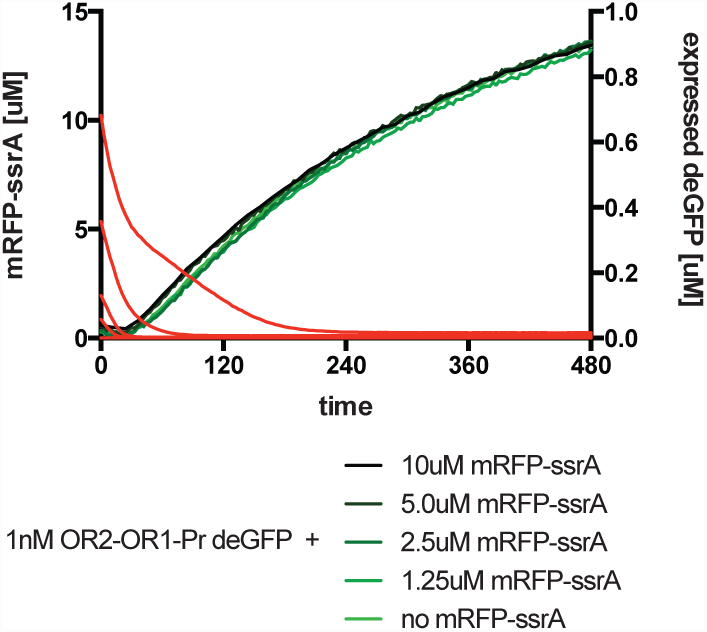
Effect of background degradation processes to TX-TL expression. A plasmid expressing deGFP off of a strong promoter-UTR is added to a TX-TL reaction at 1nM. In the same reaction, purified mRFP-ssrA is being degraded by 200 nM of added ClpX. No additional ATP or Mg is added to the reaction. 5 difference concentrations of mRFP-ssrA are tested. Both remaining mRFP-ssrA and deGFP expression are measured in real time.

We also explored increasing the amount of ClpP in a saturated ClpX TX-TL reaction to verify that ClpP was a limiting reagent. For simplicity, we expressed ClpP off of a strong promoter instead of adding purified amounts of ClpP. In the main extract used, “eZS6” derived from an ExpressIQ strain (New England Biolabs), ClpP was able to marginally increase degradation of deGFP-ssrA in the presence of 200 nM of ClpX (**Fig. 7a**). However, the ability to increase degradation was more pronounced in an alternate extract, “e8”, made from a BL21 Rosetta2 strain (Novagen) (**Fig. 7b**). This indicates that ClpXP degradation dynamics can be different depending on strain and on preparation, which could vary the amount of native ClpX or ClpP already present.

**Figure 7.**
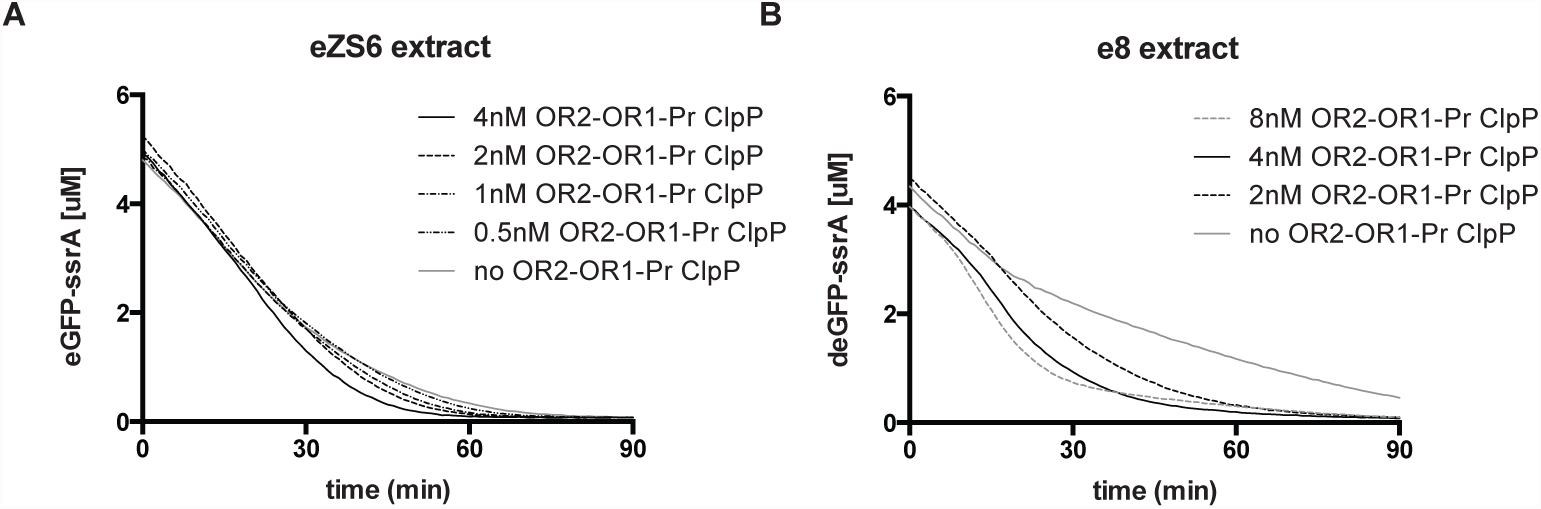
Effect of ClpP expression on degradation. A) In eZS6 extract, a plasmid expressing ClpP off of a strong promoter-UTR is added to a TX-TL reaction at varying concentrations in the presence of 200 nM of added purified ClpX. Degradation of deGFP-ssrA is plotted versus time. B) The same experiment as A), but in a different extract.

## CONCLUSION

We explored the use of supplementing the ClpXP system inherent in TX-TL with purified ClpX protease to provide fine-tunable degradation to run synthetic dynamic circuits. ClpXP degradation closely follows Michaelis-Menten kinetics, but has varying degradation rates depending on substrate. Degradation can also be controlled by ClpX concentration, allowing for up to 5-fold increased degradation rates over non-tagged variants. Degradation does not limit the running of synthetic circuits when purified ClpX is added, and rates can be further increased by adding ClpP when ClpX reaches saturated conditions between 200-400 nM.

While ClpXP provides tunable degradation, it may not be an ideal substitute for protein dilution, as degradation is neither linear nor stepwise. However, due to the fact that degradation closely follows Michaelis-Menten kinetics, degradation can be included in a “biological breadboard”-type model [20] to predict amounts needed to implement circuit dynamics. This can be easily done for fluorescent proteins, but may require indirect assays for other commonly used circuit components such as activators or repressors. However, once substrate degradation rates are characterized supplementing ClpX is easy to accomplish and fairly predictable when adjusting for concentration.

## ACKNOWLEDGEMENTS

We thank Jan Kostecki for protein purification and size exclusion chromatography assistance, Rohit Sharma for initial testing of AAA+ degradation mechanisms, Robert Sauer and Karl Schmitz for advice in purifying ClpX, and Kyle Martin for laboratory assistance. This material is based upon work supported in part by the Defense Advanced Research Projects Agency (DARPA/MTO) Living Foundries program, contract number HR0011-12-C-0065 (DARPA/CMO). Z.Z.S. is also supported by a UCLA/Caltech Medical Scientist Training Program fellowship, and a National Defense Science and Engineering Graduate fellowship. The views and conclusions contained in this document are those of the authors and should not be interpreted as representing officially policies, either expressly or implied, of the Defense Advanced Research Projects Agency or the U.S. Government.

## MATERIALS AND METHODS

### Cell-free extract preparation and execution

Preparation and execution of TX-TL was according to previously described protocols [1], with a modification in strain to ExpressIQ (New England Biolabs) and lysis according to work by Jewett et al. (personal communication). This resulted in extract “eZS6” with conditions: 9.9 mg/mL protein, 9.5 mM Mg-glutamate, 95 mM K-glutamate, 0.33 mM DTT, 1.5 mM each amino acid except leucine, 1.25 mM leucine, 50 mM HEPES, 1.5 mM ATP and GTP, 0.9 mM CTP and UTP, 0.2 mg/mL tRNA, 0.26 mM CoA, 0.33 mM NAD, 0.75 mM cAMP, 0.068 mM folinic acid, 1 mM spermidine, 30 mM 3-PGA, 2% PEG-8000. Only “eZS6” is used to prevent extract-to-extract variation, except in the final figure where “e8” prepared from BL21 Rosetta2 (Novagen) using previously described protocols [1] was used. Reactions were conducted in 15 μL in a 384-well plate (Nunc) at 29°C, and read in a Synergy H1/MF or H4 plate reader (Biotek). Settings used were: deGFP, 485 nm/515 nm gain 61; Venus, 505 nm/ 535 nm, gain 61; mRFP, 580 nm/ 610 nm, gain 100.

### Protein purification

For fluorescent proteins eGFP, mRFP, and Venus and variants eGFP-ssrA, mRFP-ssrA, and Venus-ssrA, coding sequences were cloned into a T7-lacO inducible vector containing a N-terminus His6 tag using standard techniques and propagated in a BL21-DE3 strain (New England Biolabs). Proteins were purified following a similar protocol as in [7], but were grown in TB broth in lieu of LB broth, induced with 1 mM IPTG (final concentration), and selected for a band between 25 kDa – 35 kDa corresponding to the fluorescent protein in question. Fluorescent proteins were further processed in a Supradex 20 10/300 column to select for pure, active proportions, and flash-frozen at −80°C in a storage buffer consisting of: 50 mM Tris-Cl pH 7.5, 100 mM NaCl, 1 mM DTT, 1 mM EDTA, 2% DMSO shown previously to be amenable to usage in TX-TL [7]. Final concentrations were: deGFP-ssrA, 164.8 μM; deGFP, 184.8 μM; mRFP-ssrA, 185.6 μM; mRFP, 170.6 μM; Venus-ssrA, 87.9 μM; Venus, 147.5 μM.

For ClpX, a monomeric N-terminal deletion variant Flag-clpXdeltaNLinkedHexamer-His6 was used [13] (Addgene #22143), as purifying the wildtype with a N-terminal His-tag using general Ni-NTA purification techniques resulted in a loss of activity. We followed a Ni-NTA purification procedure listed in [12], followed by Supradex 20 10/300 and functional testing for pure, active proportions above 250 kDa. Active proportions were flash frozen in the same buffer used for fluorescent proteins. Final concentration of ClpX was 1.95 μM.

